# Investigation of working memory networks for verbal and rhythmic stimuli

**DOI:** 10.1101/847038

**Authors:** Joshua D. Hoddinott, Dirk Schuit, Jessica A. Grahn

## Abstract

Auditory working memory is often conceived of as a unitary capacity, with memory for different auditory materials (syllables, pitches, rhythms) thought to rely on similar neural mechanisms. One spontaneous behavior observed in working memory studies is ‘chunking’. For example, individuals often recount digit sequences in groups, or chunks, of 3 to 4 digits, and this chunking improves performance. Chunking may also operate in musical rhythm, with beats acting as chunk boundaries for tones in rhythmic sequences. Similar to chunking, beat-based structure in rhythms also improves performance. Thus, beat processing may rely on the same mechanisms that underlie chunking of verbal material. The current fMRI study examined whether beat perception is a type of chunking, measuring brain responses to chunked and unchunked letter sequences relative to beat-based and nonbeat-based rhythmic sequences. Participants completed a sequence discrimination task, and comparisons between stimulus encoding, maintenance, and discrimination were made for both rhythmic and verbal sequences. Overall, rhythm and verbal working memory networks overlapped substantially. When comparing rhythmic and verbal conditions, rhythms activated basal ganglia, supplementary motor area, and anterior insula, compared to letter strings, during encoding and discrimination. Letter strings compared to rhythms activated bilateral auditory cortex during encoding, and parietal cortex, precuneus, and middle frontal gyri during discrimination. Importantly, there was a significant interaction in the basal ganglia during encoding: activation for beat-based rhythms was greater than for nonbeat-based rhythms, but verbal chunked and unchunked conditions did not differ. The significant interaction indicates that beat perception is not simply a case of chunking, suggesting a dissociation between beat processing and grouping mechanisms that warrants further exploration.

## 1. Introduction

The most influential model of auditory working memory is Baddeley’s phonological loop model (1). This model is largely developed on the basis of studies that use linguistic and verbal material, but may also account for auditory working memory processing of other material, such as rhythm. Verbal working memory studies indicate that grouping or ‘chunking’ is a spontaneous behavior that benefits working memory performance (2). When individuals verbally recount digit strings (such as in digit span tasks) they recount 3 or 4 numbers at a time, even when no grouping of the numbers is present at encoding. This chunking in verbal memory may be analogous to beat perception in rhythm, with beats acting as boundaries for groups of time intervals in the rhythm. Several studies have shown that temporal sequences with a regular beat are more accurately reproduced than temporal sequences with no beat (3, 4–6). Beat-based structure may enable encoding of rhythmic patterns in chunks (7), and the same neural mechanisms may therefore underlie chunking-based performance improvements in verbal working memory and beat-based rhythms. The current fMRI study examines this possibility, comparing brain responses to chunked versus unchunked verbal sequences and beat-based versus nonbeat-based rhythmic sequences.

### 1.1. Auditory working memory for verbal and rhythmic stimuli

Most models of auditory working memory posit two components: one involved in representing the to-be-remembered items and the other involved in maintaining those representations (1, 8). In Baddeley’s influential model of working memory (1), the first component is the ‘phonological short-term store’, a storage buffer for auditory memory traces. The traces decay rapidly, therefore the ‘articulatory loop’ is required to maintain items in auditory memory through active rehearsal (9). Auditory stimuli automatically enter the phonological store, which acts as an ‘inner ear’, remembering the sounds in the correct order. The articulatory process acts as an ‘inner voice’, repeating the information to prevent it from decaying. A third component, called a timing signal, was recently proposed to mark the serial order of items in the store (10, 11). However, as this component signals the order of items, not the absolute timing between items, it is unclear how the specific temporal intervals in a rhythm might be represented in the model.

Although it is unclear exactly how the model represents rhythms, previous work suggests that the articulatory loop component is important for remembering rhythms (12–14). For example, Saito (15) asked participants to listen to a short rhythm and reproduce it after a five-second delay. During encoding and delay participants either engaged in articulatory suppression (silently mouthing the vowels AEIOU) or drawing of squares. Articulatory suppression occupies the articulatory loop, whereas square drawing does not. Articulatory suppression interfered with rhythm performance much more than square drawing, suggesting that rhythm encoding and maintenance rely on the articulatory loop.

Similarly, rhythmic tapping can interfere with verbal working memory, although the interference depends on task parameters. Tapping a regular (isochronous) rhythm may interfere with short-term memory, depending on whether it is self-paced or externally paced. When tapping is self-paced, participants may simply adjust their rate of tapping to align with the presentation or rehearsal rate of the to-be-remembered verbal stimuli. However, externally paced regular tapping does interfere with verbal working memory performance when performed at a different rate from the verbal stimuli (10). Rhythmic tapping of complex rhythms interferes with verbal working memory performance regardless of whether it is self-paced or externally-paced (14).

Additional evidence for a link between auditory working memory and rhythm processing comes from behavioral measures of individual differences. Auditory working memory capacity varies across individuals, as assessed by span tasks, such as digit span. To measure digit span, a list of spoken digits is heard then repeated. The number of correctly recalled digits is the capacity of the short-term memory store, is usually between five and nine items (16). Digit span correlates with the ability to reproduce rhythms, further supporting the idea that the systems for remembering verbal and rhythmic material may overlap (17).

Finally, there is substantial neural evidence to suggest overlap between working memory and rhythm processing networks. A recent review of working memory studies (18) found consistent activation in Broca’s area, pre-supplementary motor area (pre-SMA), dorsal and ventrolateral premotor cortex (PMC), inferior frontal gyrus, cerebellum (lobule VI), as well as the intraparietal sulcus (IPS) and lateral prefrontal cortex. Subcortical activations were found in bilateral thalamus and left basal ganglia. Verbal compared to non-verbal tasks were more likely to recruit left Broca’s area, whereas non-verbal tasks were more likely to recruit left pre-SMA, SMA, and bilateral dorsal PMC. Articulatory rehearsal, specifically, activates a subset of these working memory areas, including Broca’s area, SMA and pre-SMA, dorsal and ventrolateral PMC, cerebellum, and anterior insula (19–23). These areas are also commonly activated in studies of rhythm perception and production (4, 24–26), suggesting reliance on at least partially overlapping neural processes.

Previous work has specifically compared working memory for verbal and musical material, but has generally focused on pitch, not rhythm (27, 28). Consistent with the review study above (18), these studies find overlap between the brain areas involved in working memory for verbal and pitch sequences. Activations are observed in parietal cortex (supramarginal gyrus (SMG) and intraparietal sulcus (IPS)), posterior temporal cortex (planum temporale or area SPt), ventrolateral and dorsolateral PMC, Broca’s area, and dorsolateral cerebellum (27–29). More recent studies (30, 31) find similar overlap between verbal and pitch stimuli during rehearsal: vlPMC and dlPMC, the anterior insula, the SMG/IPS, the planum temporale, the IFG, pre-SMA, basal ganglia, and the cerebellum. The authors suggest that vlPMC and Broca’s area are part of an active rehearsal component of the articulatory loop, as these areas did not respond to simply subvocalizing without a rehearsal function. Therefore, these areas might be expected to respond during rehearsal of rhythm as well.

### 1.2. Beat Perception and Chunking

Both beat perception (in rhythm) and chunking (in verbal sequences) are known to improve working memory performance. Beat perception spontaneously arises in the context of auditory rhythm: humans perceive a regular pulse, or beat, that marks equally spaced points in time (32, 33). Perception of a beat occurs without effort as long as the auditory sequence has a regular temporal structure, such as periodically occurring event onsets in the range of ~300-900 ms (34, 35). Several studies confirm that beat perception leads to higher accuracy in rhythm synchronization, discrimination, and reproduction (3, 5, 6, 36).

Chunking refers to the process of taking individual units of information and grouping them into larger units (chunks). A common example of chunking occurs in telephone phone numbers, in which individual digits are grouped into 3- and 4-unit chunks. Chunking is a useful method for information reduction. By grouping individual elements into larger blocks, information becomes easier to retain and recall. The grouping of sequential units into a single, larger, unit to facilitate performance is observed in a variety of domains (37, 38). For example, many chunking studies have been conducted in the motor learning domain, and find that sequences of finger movements are spontaneously chunked (39–41), reducing working memory load during ongoing performance (37, 42).

### 1.3. Chunking and Beat perception in fMRI

Chunking has been investigated using fMRI. One previous study presented 6-letter visual sequences (43), and found that chunked sequences (a brief pause was inserted between 3-letter groups) evoked greater activity in right inferior frontal gyrus (BA 47). However, the findings may not readily apply to the auditory modality, as chunking effects are generally smaller for visual compared to auditory sequences (44–46). More recently, Kalm et al. (47) investigated chunking using auditory sequences of 6 or 9 letters. Chunking resulted in reduced activity in auditory areas. For 9-letter strings only, parietal areas were more active for chunked than unchunked strings. When comparing activation differences for chunked and unchunked letter strings to activation differences observed between beat and nonbeat rhythms, not much overlap exists, as beat compared to nonbeat rhythms activate the basal ganglia, SMA, and sometimes the left inferior frontal gyrus (6, 48).

In the current study we tested whether beat perception and chunking rely on similar neural mechanisms, using a working memory paradigm that could be applied to both verbal and rhythmic stimuli. Each trial consisted of stimulus presentation (encoding), a silent delay for rehearsal (maintenance), and a second stimulus presentation (discrimination), after which participants indicated whether the second stimulus was the same as or different from the first. Half of the trials involved discriminating rhythmic sequences (comparing two rhythms to determine whether the timing was same or different), and the other half involved discriminating letter sequences (comparing two strings of different letters to determine whether letter order was the same or different). Half of the rhythm trials used beat-based sequences, which induced a perception of a regular beat, and the other half were nonbeat-based sequences, with irregular timing in which no beat perception was possible. Similarly, on half of the letter trials, the timing of the letter presentation created two equal chunks, and on the other half, the timing was irregular, with no chunks.

## 2. Methods

### 2.1. Participants

18 volunteers (4 female; *M*_age_ = 28.3, *SD* = 8.65) participated in the brain imaging study. All participants completed the experiment and received financial compensation for participation. The Cambridge University Psychological Research Ethics Committee provided clearance for the study (CPREC 2009.17).

### 2.2. Stimuli

Recordings of spoken letters from a single male speaker were used to construct all stimuli. The stimuli for the rhythm reproduction task were created using GarageBand (Apple, Inc., v4.1.2 (248.7)). Beat rhythms were constructed using the following six core patterns: 1111, 112, 211, 22, 31, 4, similar to previous work (6, 49). Short and medium rhythms consisted of two or three core patterns (e.g., 1122114), respectively. The shortest interval (i.e., 1) ranged from 220-280 milliseconds, in 10 millisecond steps, creating seven potential tempi. The other intervals in the rhythm were integer multiples of the shortest interval. One final note was added to the end of each sequence to mark the end of the last interval. None of the six core patterns were repeated within a rhythm. On each trial, one of the seven different tempi was used. The trial-to-trial tempo change prevented carry-over of the beat from one trial to the next trial.

Beat rhythms were modified to create nonbeat versions. One third of the intervals in each rhythm kept their original length, one third were increased in length by 1/3 of 1 unit and one third decreased in length by 1/3 of 1 unit. Thus, the nonbeat rhythms were the same as the beat rhythms in overall duration and number of intervals, but had irregular timing (see Figure 1).

**Figure 1.**
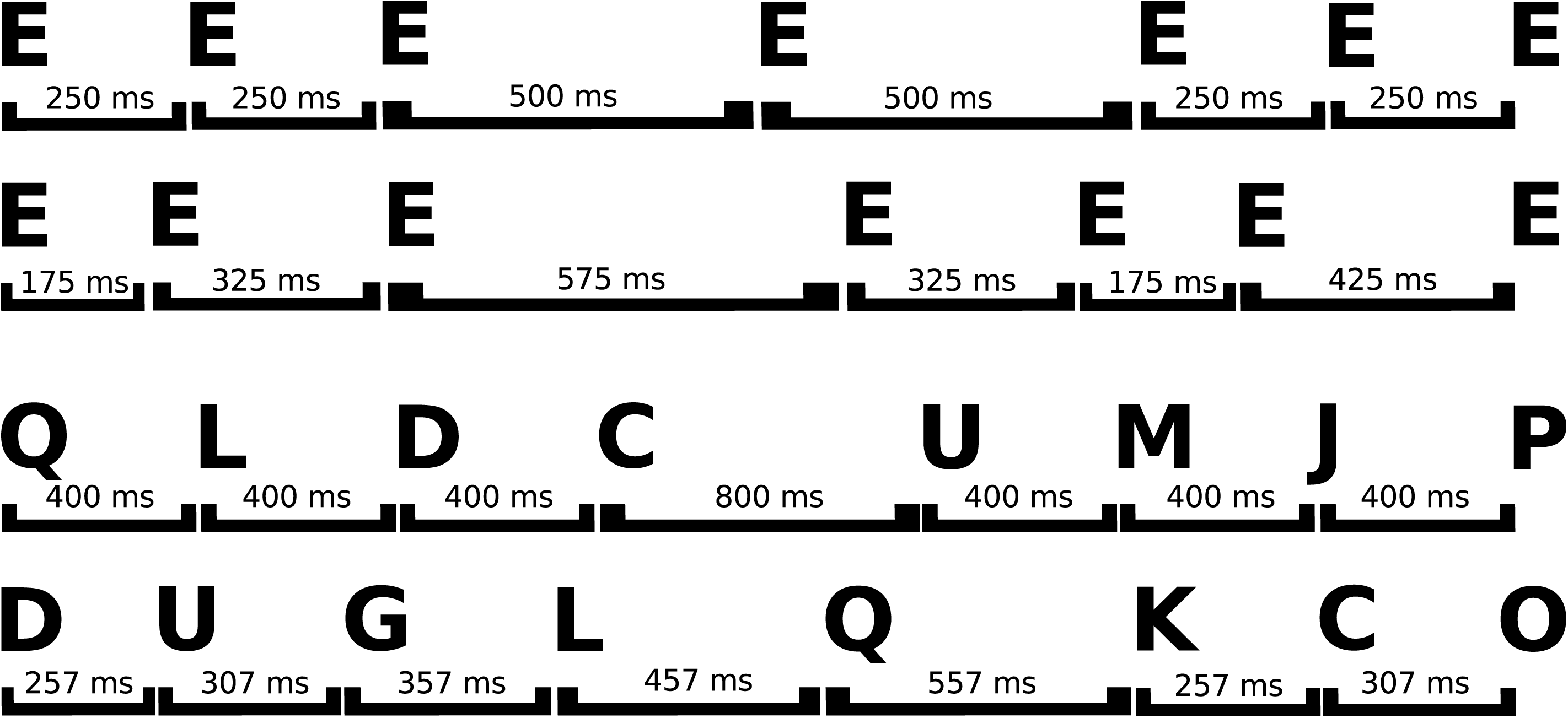
Schematic examples of the four different stimulus types, constructed with spoken letters. Time intervals between letter onsets are given in milliseconds (ms). A) A Beat rhythm (upper) and nonbeat rhythm (lower). B) A chunked verbal sequence (upper) and an unchunked verbal sequence (lower).

For the letter sequences, strings of four (short length) or eight (medium length) different, easily intelligible, letters were created (e.g., ‘Q L D C U M J P’). Half of the sequences had regular timing and half had irregular timing. For the regular sequences, the strings were divided in half (two groups of two letters, or two groups of four letters), ensuring that the sequences were encoded as chunks (50, 51). The letter onsets within a group were separated by 400 ms, and each group was separated by 800 ms. The letter onsets in the irregular strings were separated by unequal time intervals. The four-letter strings used 233, 533 and 833 ms intervals (in random order). The eight-letter strings used 257, 307, 357, 457, 557 and 657 ms intervals (again, in random order). Sample strings are shown in Figure 1.

### 2.3. Design

A 2 × 2 within-subject design was used, with experimental factors *stimulus type* (rhythm, letter) and *temporal regularity* (beat/nonbeat for rhythms, chunked/unchunked for letters) for each stage (encoding, maintenance, discrimination of same rhythms, and discrimination of different rhythms). There were three trial types: full trials, stimulus-response-only trials and null trials. Full trials consisted of encoding, maintenance, and discrimination periods as well as a response. For the participants, each period was distinguished by a differently coloured display. During the *encoding period*, the initial stimulus was heard with a blue display. The subsequent *maintenance period* had a black display and lasted zero, one, or two times the length of the preceding stimulus. The *discrimination period* stimulus was heard with a green display. The stimulus was either the same as or different from the first stimulus. For ‘different’ stimuli, sequence of the same type as the stimulus (beat/chunked or nonbeat/unchunked) was used. During the *response period*, the screen was red and participants had two seconds to indicate whether the stimuli were same or different with a button press. Stimulus-response-only trials had only the initial stimulus presentation, after which the screen turned red and text instructed the participant to press the left or right button. Null trials consisted of a 9-second blank screen. The variable length of maintenance periods, stimulus-response-only trials, and null trials were necessary to allow the hemodynamic response to the different trial periods to be de-correlated and therefore estimable. The variable delay decouples the encoding period from the maintenance period and also the maintenance period from the discrimination period. The stimulus-response-only trials decouple the encoding period from the discrimination period and the discrimination period from the response.

There were 16 blocks in the experiment. One block comprised eight full trials (2 with zero length maintenance, 4 with 1x length, 2 with 2x length), two stimulus-response-only trials and two null trials. Each block contained only rhythm or letter trials. The stimulus type alternated (rhythm, letter condition) with each block.

### 2.4. Procedure

Participants gave written informed consent and practiced one rhythm and one letter block prior to entering the scanner. Each practice block contained four full trials and one stimulus-response-only trial. All stimuli were unique to the practice session.

### 2.5. MR scanning specifications

A 3T Siemens Tim Trio MRI scanner was used to collect two runs with 540 echoplanar imaging (EPI) volumes in each. All EPI data had 36 slices, matrix size of 64 × 64, echo time (TE) = 30 ms, repetition time (TR) = 2.19 s, field of view = 19.2 × 19.2 cm, flip angle = 78 degrees, slice thickness 3 mm, interslice distance of 0.75 mm, and in-plane resolution of 3 × 3 mm. High-resolution MPRAGE anatomical images (TR = 2250 ms, TE = 2.99 ms, flip angle = 9 degrees, inversion time = 900 ms, 256 × 256 × 192 isotropic 1 mm voxels) were collected for anatomic localization and coregistration.

### 2.6. Data pre-processing and analysis

SPM8 was used for data analysis (SPM8; Wellcome Centre for Neuroimaging, London, UK). The first five EPI volumes of each run were discarded to allow for T1 equilibration. Images were sinc-interpolated in time to correct for acquisition time differences within each volume and realigned spatially with respect to the first image of the first run using trilinear interpolation. The coregistered MPRAGE image was segmented and normalized using affine and smoothly nonlinear transformations to the T1 template in Montreal Neurological Institute (MNI) space. The normalization parameters were then applied to the EPIs and all normalized EPI images were spatially smoothed with a Gaussian kernel of full-width half-maximum 8 mm. For each participant, encoding, maintenance, discrimination, and response were modelled separately for each condition. These were modelled using a regressor made from an on-off boxcar convolved with a canonical hemodynamic response function (apart from response, which was modelled with a delta function convolved with the canonical hemodynamic response function). EPI volumes associated with discrete artifacts were included as covariates of no interest (nulling regressors). This included volume displacements >4mm or spikes of high variance in which scaled volume to volume variance was 4 times greater than the mean variance of the run. Autocorrelations were modelled using an AR(1) process and low-frequency noise was removed with a standard high-pass filter of 128 seconds.

The contrast images estimated from single participant models were entered into second-level random effects analyses for group inference (52). Separate ANOVAs were conducted for encoding, maintenance, same discrimination, and different discrimination periods. Each ANOVA was 2 × 2 with the factors *temporal regularity* and *stimulus type*. All effects were estimated using t-contrasts. Significance level was α = .05, using cluster-wise False Discovery Rate correction (53), and cluster-forming threshold of p <.001 uncorrected.

In addition, for each stage (encoding, maintenance, discrimination same, discrimination different) regions of interest for the average of all conditions versus rest were created. Each region was defined as a 10-mm radius sphere around the peak voxel in a region, except putamen which was 5 mm. ANOVAs (2 × 2, as above) were conducted on each region’s activity. This enabled us to test more sensitively for differences between conditions, using orthogonally-defined task-relevant regions. The specific regions tested are given in the Supplementary material. Effects are reported here for brain regions not already identified as significant by whole-brain analyses. Significance level was α = .05, Bonferroni-corrected for number of regions tested at that particular stage. Thus, Encoding: α =.0031; Maintenance: α = .0045; Discrimination same: α =.0033; Discrimination different: α = .0031.

## 3. Results

### 3.1. Behavioral results

Performance accuracy (percentage of correctly discriminated sequences) was compared for each condition. Overall, accuracy was high (*M* = 86%, *SD* = 6.3%), but better in the verbal conditions than in the rhythm conditions (*M*_rhythm beat_ = 83%, *SD* = 11%; *M*_rhythm nonbeat_ = 73%, *SD* = 12%; *M*_verbal chunked_ = 94%, *SD* = 6%; *M*_verbal unchunked_ = 93%, *SD* = 4%). A 2 (stimulus type) x 2 (temporal regularity) repeated measures ANOVA on percent correct scores confirmed a significant interaction (*F*(1, 17) = 22.39, *p* < .001, η_p_^2^ = .57), driven by a significant difference in performance between beat and nonbeat rhythms (*t*(1, 17) = 5.13, *p* < .001) but no significant difference between verbal chunked and unchunked conditions (*t*(1,17) = 1.1, *p* = .27). As the behavioral performance differed across conditions, only correct trials were modeled in the fMRI analyses.

### 3.2. fMRI results: whole brain analyses

#### 3.2.1. Encoding

Several brain areas responded to the encoding stage, across all conditions. These include bilateral superior temporal gyri, premotor cortex, SMA, inferior frontal gyrus, parietal cortex, cerebellum, and basal ganglia (see Figure 2 and Supplementary table 1 for coordinates of maxima). Contrast analysis across verbal and non-verbal conditions (Verbal > Non Verbal) shows activation in bilateral superior and middle temporal gyri for letter strings more than rhythms (see Supplementary Table 2), likely because of the greater acoustic variety in letter strings compared to rhythms. In the opposite contrast (Non Verbal > Verbal), rhythms activated the basal ganglia, SMA, left cerebellum and right inferior frontal gyrus more than letter strings (Table 1). A significant interaction between stimulus type and temporal regularity was found in the bilateral basal ganglia (see Tables 1–3). As shown in Figure 2, this stems from greater activation for beat than nonbeat rhythms, but no difference between chunked and unchunked letter strings.

**Table 1.** Stereotaxic locations of peak voxels during encoding: rhythm condition – verbal condition.

| Brain area | Brodmann area | Cluster* | voxel t | pFDR (cluster) | x | y | z |
| --- | --- | --- | --- | --- | --- | --- | --- |
| L inferior frontal gyrus, p. triangularis | BA 47 | cluster 3 | 3.38 | <.001 | -48 | 21 | 0 |
| L inferior frontal gyrus, p. triangularis | BA 45 | cluster 3 | 3.92 | <.001 | -57 | 21 | 3 |
| R inferior frontal gyrus, p. triangularis | BA 45 | cluster 6 | 4.23 | <.05 | 42 | 30 | 9 |
| R inferior frontal gyrus, p. triangularis | BA 47 | cluster 6 | 3.65 | <.05 | 36 | 30 | 3 |
| L inferior frontal gyrus, p. opercularis | BA 6 | cluster 3 | 4.13 | <.001 | -48 | 9 | 15 |
| R inferior frontal gyrus p. opercularis | BA 44 | cluster 1 | 5.89 | <.001 | 54 | 12 | 24 |
| R inferior frontal gyrus p. opercularis | BA 6 | cluster 1 | 5.02 | <.001 | 54 | 9 | 15 |
| L insula |  | cluster 3 | 4.07 | <.001 | -36 | 3 | -3 |
| L insula |  | cluster 3 | 4.46 | <.001 | -45 | 0 | 6 |
| L insula |  | cluster 3 | 4.38 | <.001 | -39 | 0 | 6 |
| L supplementary motor area | BA 6 | cluster 4 | 4.32 | <.01 | -9 | -3 | 66 |
| R supplementary motor area | BA 6 | cluster 4 | 4.24 | <.01 | 6 | 3 | 66 |
| R premotor cortex | BA 6 | cluster 1 | 3.46 | <.001 | 48 | 3 | 42 |
| L Rolandic operculum |  | cluster 3 | 4.58 | <.001 | -51 | 3 | 12 |
| R middle temporal gyrus | BA 21 | cluster 5 | 4.24 | <.05 | 48 | -36 | 51 |
| L amygdala | BA 36 | cluster 3 | 4.31 | <.001 | -27 | 0 | -24 |
| L pallidum |  | cluster 3 | 4.57 | <.001 | -21 | 6 | 3 |
| L pallidum |  | cluster 3 | 4.36 | <.001 | -15 | 0 | -9 |
| R pallidum |  | cluster 2 | 5.70 | <.001 | 18 | 3 | -6 |
| L putamen |  | cluster 3 | 4.64 | <.001 | -18 | 12 | 3 |
| R putamen |  | cluster 2 | 5.48 | <.001 | 21 | 3 | 15 |

Cluster volumes: 1=322 voxels, 2=286 voxels, 3=328 voxels, 4=96 voxels, 5=61 voxels, 6=56 voxels

**Table 2.** Stereotaxic locations of peak voxels during encoding: Interaction of (beat rhythm condition – nonbeat rhythm condition) – (chunked letter sequences – unchunked letter sequences)

| Brain area | Cluster* | voxel t | pFDR (cluster) | x | y | z |
| --- | --- | --- | --- | --- | --- | --- |
| L insula | cluster 2 | 3.82 | <.01 | -27 | 18 | -6 |
| R pallidum | cluster 2 | 3.85 | <.1 | 18 | 3 | -3 |
| L putamen | cluster 1 | 4.74 | <.01 | -24 | 9 | 3 |

Cluster volumes: 1=121 voxels, 2=46 voxels

**Table 3.** Stereotaxic locations of peak voxels during encoding: beat rhythm condition – nonbeat rhythm condition.

| Brain area | Cluster* | voxel t | pFDR (cluster) | x | y | z |
| --- | --- | --- | --- | --- | --- | --- |
| L putamen | cluster 1 | 5.87 | <.005 | -24 | 3 | 0 |
| R putamen | cluster 2 | 5.18 | <.005 | 21 | 9 | 6 |
| R putamen | cluster 2 | 4.39 | <.005 | 24 | 6 | -3 |

Cluster volumes: 1=178 voxels, 2=140 voxels

**Figure 2.**
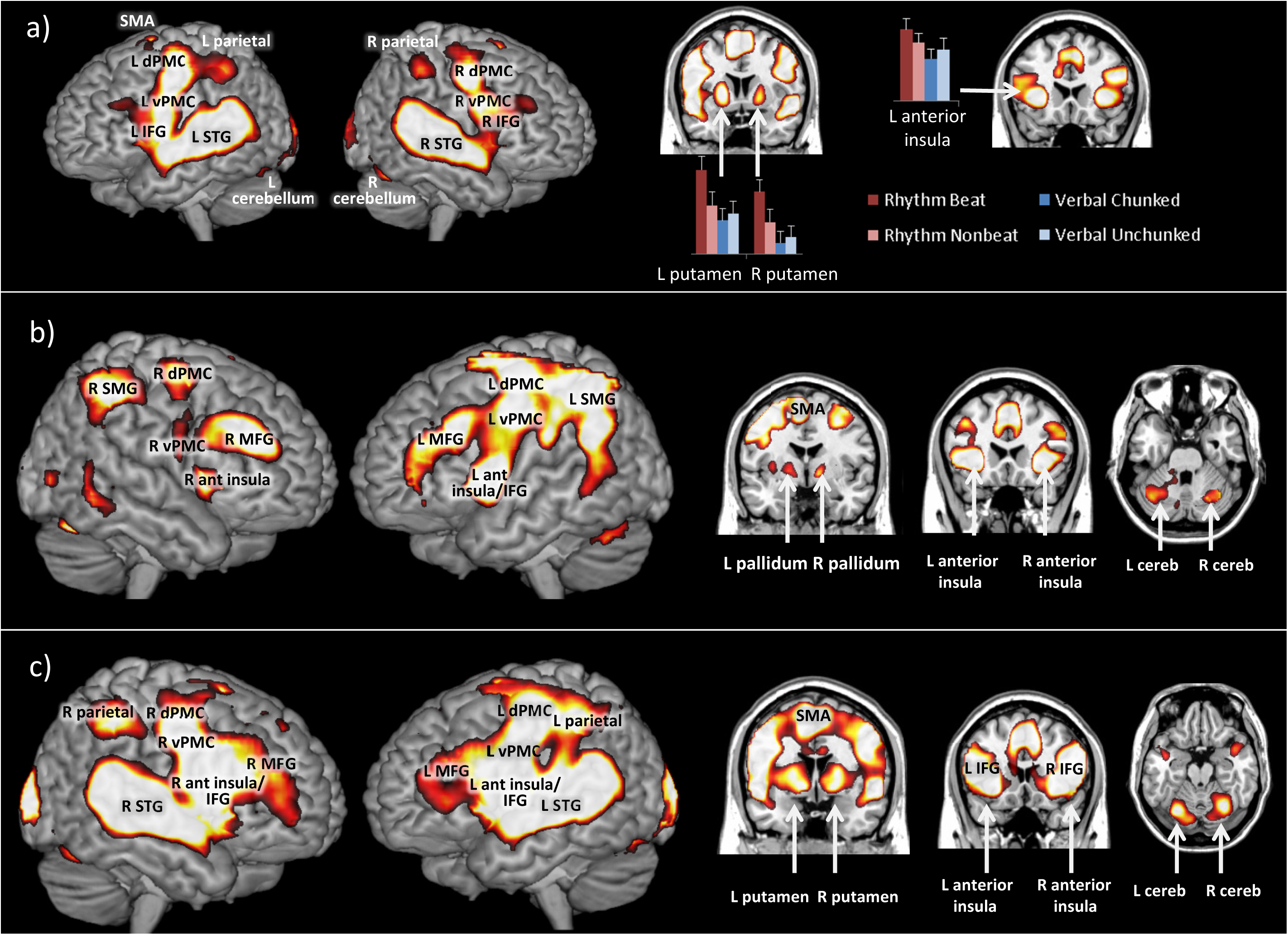
Significant activation for encoding (a), maintenance (b), and discrimination phases (c). The brain renderings show whole brain contrasts for all conditions (beat rhythms, nonbeat rhythms, chunked letter sequences, and unchunked letter sequences) versus rest. Inset Bar graphs show activation levels for each condition in brain regions that showed significant interactions between conditions. (L, left; R, right; cereb, cerebellum; dPMC, dorsal premotor cortex; vPMC, ventral premotor cortex; IFG, inferior frontal gyrus; MFG, middle frontal gyrus; SMG, supramarginal gyrus; STG, superior temporal gyrus; SMA, supplementary motor area.)

#### 3.2.2. Maintenance

Across all conditions, activation was observed in bilateral SMA, premotor cortex, insula, inferior frontal gyri, inferior parietal cortex, and right cerebellum (see Figure 2 and Supplementary table 3). There were no other significant main effects or interactions.

#### 3.2.3. Discrimination of same sequences

Across all conditions, activation was observed in bilateral SMA, premotor cortex, insula, inferior frontal gyri, inferior parietal cortex, and superior temporal gyri. Verbal sequences elicited greater bilateral superior and middle temporal gyri, and left inferior frontal gyrus activity than rhythm sequences (see Figure 2 and Supplementary table 4). Letter strings activated bilateral superior and middle temporal gyri more than rhythms (see Supplementary Table 5). There were no other significant main effects or interactions.

#### 3.2.4. Discrimination of different sequences

The activation loci for this contrast are reported in Supplementary tables 6-7 for completeness, but this stage is associated with detection of acoustic changes and response preparation, therefore it is likely to encompass many cognitive processes that are not of interest.

### 3.3. fMRI analyses: region of interest analyses

Most ROIs that showed significant main effects or interactions in ROI analyses were already identified by whole brain analyses, with the exception of the right putamen, which showed a significant interaction between temporal regularity and stimulus type (t(1,17) = 4.14, p = .001). As shown in Figure 2, this stems from greater activation for beat than nonbeat rhythms, but no difference between chunked and unchunked letter strings. A full list of regions is shown in Supplementary table 8).

## 4. Discussion

The goal of the study was to compare the neural networks involved in working memory for verbal and rhythmic stimuli. In general, similar to previous working memory studies, the networks were characterized by overlap, rather than differences. During encoding, all conditions resulted in activation in frontoparietal areas, premotor and auditory cortices, and the basal ganglia and cerebellum. Activity in auditory areas was greater for verbal than rhythmic stimuli, and activity in motor areas was greater for rhythm than verbal stimuli. In the basal ganglia, greater activity was observed for beat than nonbeat rhythms, but no difference was found for chunked and unchunked verbal stimuli. During maintenance, activity was observed in similar areas to encoding, apart from auditory cortex. During discrimination, activity was observed in similar areas to encoding, but generally with greater spatial extent. Greater auditory activity was observed for verbal than rhythmic stimuli. For maintenance and discrimination, no interactions between stimulus type and temporal structure were observed. The activity associated with the working memory task is consistent with previous research that suggests a ‘core’ working memory network that includes inferior frontal areas, parietal areas, pre-SMA, PMC, cerebellum, left basal ganglia (18).

Activation differences between rhythm and verbal stimuli were mainly observed during the encoding and discrimination stages. In both stages, verbal stimuli compared to rhythmic stimuli elicited greater activity in the auditory cortex. However, an absence of interactions with temporal regularity in the auditory cortex activation suggests that observed auditory activation differences are driven by basic stimulus differences. This is likely because of the acoustic variation associated with hearing a string of different letters during verbal stimuli, instead of a repetition of single letters during rhythm stimuli.

During encoding only, rhythmic stimuli elicited greater activity (relative to verbal stimuli) in the basal ganglia (putamen and pallidum) as well as the SMA and bilateral inferior frontal cortex. This is broadly consistent with the idea that motor areas may play a greater role in the processing of rhythmic auditory information than in processing auditory identity. A meta-analysis (18) contrasted verbal (e.g., letters, words) and non-verbal (e.g., figures, objects) stimuli. They found left Brodmann areas 44 and 45 are more closely associated with verbal stimuli. By their criteria, both the rhythm and letter conditions would be ‘verbal’, and indeed both rhythm and letter conditions activate this area. However, we find even greater activity for rhythm than verbal stimuli. Therefore, the focus on the temporal aspects of the stimuli as opposed to the identities of the letters appears to recruit IFG, perhaps because of its role in temporal sequencing (54, 55).

During encoding an interaction was observed in the basal ganglia bilaterally, with beat rhythms inducing significantly greater activity than nonbeat rhythms, and also than chunked and unchunked verbal stimuli. Thus, the effects of temporal structure appear to be most influential during the encoding of stimuli, rather than during rehearsal (the maintenance period) and discrimination. The presence of a significant interaction in the basal ganglia suggests that beat perception is not simply a form of chunking, and that beat perception recruits the basal ganglia specifically. This is consistent with other neuroimaging work that associates the basal ganglia with temporal structure, particularly in discrimination tasks (6, 56, 57). The presence of an interaction during encoding but not maintenance or discrimination suggests a few possibilities about the role of the basal ganglia in beat perception, although it must also be emphasized that any interpretation of null results must be treated with caution. It may be that the basal ganglia are important for an initial reorganization of rhythmic stimuli that occurs when a beat is perceived—intervals are recoded from a series of separate durations to onsets relative to the beat. The motivation to recode the intervals is high in a discrimination task, when performance depends on an accurate representation, and beat perception can enhance performance. It is also possible, contrary to the conclusions of previous research (58), that the basal ganglia are important for the perception of the beat, rather than maintenance or internal generation of the beat during the maintenance period. We feel that this interpretation, though possible, is less likely, as subthreshold differences in beat and nonbeat rhythms were observed during the maintenance period.

We did not find any clear differences between verbal chunked and unchunked stimuli. Previous work has reported greater right IFG activation for chunked than unchunked sequences, although the sequences were visual (10). Kalm et al. (47) found differences in parietal cortex activity for chunked compared to unchunked auditory letter sequences, but only when the sequences were nine letters long, not six, and therefore exceeded auditory working memory span. Our sequences were four and eight letters long, so may not have been long enough to reliably exceed our participants’ spans. We assessed behavioral performance with discrimination, not recall, which is usually used to determine span. However, performance was very high in both the chunked and unchunked conditions (>90%), therefore the task may not have been taxing enough to elicit neural activation differences between chunked and unchunked conditions.

Finally, although we did not observe similarities in neural activation to beat perception and chunking, we only tested a perceptual task. In contrast to the neural areas for chunking of perceptual sequences tested by previous work, a role for the basal ganglia in chunking has been proposed in the motor sequencing literature. The basal ganglia may link sequential responses together into chunks. Basal ganglia neurons appear to represent chunks by firing preferentially at the beginning and end of action sequences (59–61), and disruption of this firing impairs sequence learning (62). Moreover, individuals with basal ganglia dysfunction, such as in Parkinson’s disease (63) or stroke (64), are less likely to chunk motor sequences. Therefore, the motor sequencing literature suggests that there may be neural overlap between chunking of movement sequences and beat perception in auditory sequences, namely in the basal ganglia. Further research to compare chunking in motor sequences and beat perception may support the idea that beat perception engages some fundamentally motor aspects, mediated by auditory-motor interactions.

## Acknowledgements

Thank you to Matt Davis for assistance and advice about stimulus creation.

