## Supplementary Tables for "Investigation of working memory networks for verbal and rhythmic stimuli"

**Supplementary table 1**

**Stereotaxic locations of peak voxels during encoding: all conditions – baseline**

| Brain area | Brodmann area | Cluster* | voxel t | pFDR (cluster) | x | y | z |
|---|---|---|---|---|---|---|---|
| L inferior frontal gyrus, p. opercularis | BA 44 | cluster 1 | 31.78 | <.001 | -51 | 9 | 6 |
| L inferior frontal gyrus, p. opercularis | BA 44 | cluster 1 | 29.39 | <.001 | -51 | 9 | 21 |
| R inferior frontal gyrus, p. opercularis | BA 44 | cluster 2 | 25.45 | <.001 | 42 | 12 | 27 |
| R inferior frontal gyrus, p. opercularis | BA 44 | cluster 2 | 26.55 | <.001 | 45 | 6 | 27 |
| L anterior cingulate | BA 32 | cluster 1 | 11.27 | <.001 | -12 | 18 | 30 |
| L anterior insula | BA 47 | cluster 1 | 23.19 | <.001 | -36 | 21 | 3 |
| R anterior insula | BA 47 | cluster 2 | 22.75 | <.001 | 33 | 21 | 3 |
| L middle frontal gyrus | BA 46 | cluster 1 | 8.85 | <.001 | -33 | 39 | 24 |
| L middle frontal gyrus | BA 46 | cluster 1 | 7.93 | <.001 | -30 | 42 | 21 |
| R middle frontal gyrus | BA 46 | cluster 2 | 8.13 | <.001 | 33 | 39 | 27 |
| L supplementary motor area | BA 6 | cluster 1 | 65.93 | <.001 | -6 | 3 | 60 |
| L ventral premotor cortex | BA 6 | cluster 1 | 29.63 | <.001 | -48 | 9 | 15 |
| L premotor cortex | BA 6 | cluster 1 | 20.50 | <.001 | -30 | -6 | 51 |
| L premotor cortex | BA 6 | cluster 1 | 45.47 | <.001 | -51 | 0 | 45 |
| R premotor cortex | BA 6 | cluster 2 | 35.53 | <.001 | 51 | 3 | 45 |
| R premotor cortex | BA 6 | cluster 2 | 34.74 | <.001 | 54 | 0 | 48 |
| R premotor cortex | BA 6 | cluster 2 | 11.36 | <.001 | 30 | -3 | 45 |
| L postcentral gyrus | BA 4 | cluster 1 | 10.46 | <.001 | -39 | -21 | 54 |
| R postcentral gyrus | BA 2 | cluster 3 | 13.51 | <.001 | 48 | -33 | 48 |
| L inferior parietal lobule | BA 7 | cluster 1 | 16.65 | <.001 | -30 | -51 | 45 |
| L inferior parietal lobule | BA 40 | cluster 1 | 16.46 | <.001 | -42 | -42 | 48 |
| R inferior parietal lobule | BA 40 | cluster 3 | 20.06 | <.001 | 33 | -51 | 45 |
| L superior temporal gyrus | BA 22 | cluster 1 | 75.41 | <.001 | -66 | -21 | 6 |
| L superior temporal gyrus | BA 22 | cluster 1 | 66.67 | <.001 | -54 | -21 | 6 |
| L superior temporal gyrus | BA 41 | cluster 1 | 55.14 | <.001 | -42 | -36 | 12 |
| L superior temporal gyrus | BA 41 | cluster 1 | 53.74 | <.001 | -45 | -33 | 9 |
| R superior temporal gyrus | BA 21 | cluster 2 | 74.33 | <.001 | 63 | -30 | 6 |
| R superior temporal gyrus | BA 22 | cluster 2 | 73.43 | <.001 | 57 | -18 | 0 |
| L middle temporal gyrus | BA 21 | cluster 1 | 66.98 | <.001 | -63 | -30 | 3 |
| R middle temporal gyrus | BA 42 | cluster 2 | 53.25 | <.001 | 51 | -39 | 9 |
| L inferior temporal gyrus | BA 37 | cluster 1 | 7.96 | <.001 | -42 | -57 | -9 |
| L superior occipital gyrus | BA 17 | cluster 1 | 39.58 | <.001 | -12 | -90 | 3 |
| L fusiform area | BA 19 | cluster 1 | 13.96 | <.001 | -27 | -69 | -12 |
| R fusiform area | BA 19 | cluster 1 | 16.80 | <.001 | 30 | -75 | -6 |
| L hippocampus |  | cluster 1 | 10.50 | <.001 | -21 | -27 | -3 |
| L lingual gyrus | BA 19 | cluster 1 | 7.89 | <.001 | -15 | -51 | -3 |
| R lingual gyrus | BA 19 | cluster 1 | 7.91 | <.001 | 18 | -51 | -6 |
| R calcarine sulcus | BA 17 | cluster 1 | 48.45 | <.001 | 12 | -96 | 3 |
| L calcarine sulcus | BA 17 | cluster 1 | 39.28 | <.001 | -9 | -93 | -3 |
| R pallidum |  | cluster 2 | 11.93 | <.001 | 21 | 6 | 0 |
| L cerebellum, lobule VI |  | cluster 1 | 17.41 | <.001 | -27 | -63 | -21 |
| R cerebellum, lobule VI |  | cluster 1 | 23.17 | <.001 | 30 | -63 | -21 |

Cluster volumes: 1=10051 voxels, 2=3629 voxels, 3=459 voxels

**Supplementary table 2**

**Stereotaxic locations of peak voxels during encoding: verbal condition – rhythm condition**

| Brain area | Brodmann area | Cluster* | voxel t | pFDR (cluster) | x | y | z |
|---|---|---|---|---|---|---|---|
| R superior temporal gyrus | BA 22 | cluster 1 | 9.86 | <.001 | 57 | -18 | 0 |
| L middle temporal gyrus | BA 22 | cluster 2 | 9.38 | <.001 | -66 | -21 | 3 |
| L middle temporal gyrus | BA 22 | cluster 2 | 9.04 | <.001 | -60 | -12 | 0 |

Cluster volumes: 1=394 voxels, 2=400 voxels

**Supplementary table 3**

**Stereotaxic locations of peak voxels during maintenance: all conditions – baseline**

| Brain area | Brodmann area | Cluster* | voxel t | pFDR (cluster) | x | y | z |
|---|---|---|---|---|---|---|---|
| L middle frontal gyrus | BA 9 | cluster 1 | 13.13 | <.001 | -48 | 15 | 3 |
| L middle frontal gyrus | BA 46 | cluster 1 | 8.44 | <.001 | -27 | 42 | 21 |
| R middle frontal gyrus | BA 46 | cluster 3 | 10.64 | <.001 | 36 | 39 | 24 |
| R middle frontal gyrus | BA 46 | cluster 3 | 10.02 | <.001 | 36 | 36 | 30 |
| L inferior frontal gyrus, p. triangularis | BA 46 | cluster 1 | 10.38 | <.001 | -39 | 36 | 27 |
| L inferior frontal gyrus, p. triangularis | BA 45 | cluster 1 | 10.41 | <.001 | -42 | 30 | 30 |
| L inferior frontal gyrus, p. opercularis | BA 44 | cluster 1 | 8.69 | <.001 | -42 | 6 | 27 |
| L rostral superior frontal gyrus | BA 47 | cluster 1 | 6.41 | <.001 | -33 | 54 | 0 |
| L rostral superior frontal gyrus | BA 11 | cluster 1 | 6.66 | <.001 | -27 | 51 | -3 |
| L medial orbitofrontal gyrus | BA 47 | cluster 1 | 7.19 | <.001 | -30 | 48 | -9 |
| L anterior insula | BA 47 | cluster 1 | 13.55 | <.001 | -30 | 24 | 0 |
| R anterior insula | BA 47 | cluster 3 | 12.14 | <.001 | 33 | 24 | -3 |
| L insula | | cluster 1 | 13.18 | <.001 | -42 | 15 | 6 |
| R precentral gyrus | BA 44 | cluster 3 | 4.01 | <.001 | 51 | 9 | 39 |
| L supplementary motor area | BA 6 | cluster 1 | 15.31 | <.001 | -3 | 6 | 54 |
| L supplementary motor area | BA 6 | cluster 1 | 15.28 | <.001 | -6 | 12 | 48 |
| L premotor cortex | BA 6 | cluster 1 | 13.25 | <.001 | -33 | -6 | 63 |
| L premotor cortex | BA 6 | cluster 1 | 8.33 | <.001 | -21 | 3 | 69 |
| R premotor cortex | BA 6 | cluster 3 | 4.98 | <.001 | 63 | 9 | 18 |
| R premotor cortex | BA 6 | cluster 3 | 4.48 | <.001 | 60 | 9 | 33 |
| R premotor cortex | BA 6 | cluster 3 | 5.06 | <.001 | 63 | 9 | 27 |
| L ventral premotor cortex | BA 6 | cluster 1 | 12.42 | <.001 | -54 | 0 | 39 |
| R ventral premotor cortex | BA 6 | cluster 4 | 7.75 | <.001 | 33 | 0 | 48 |
| R ventral premotor cortex | BA 6 | cluster 4 | 7.17 | <.001 | 36 | -3 | 60 |
| L postcentral gyrus | BA 3 | cluster 1 | 8.11 | <.001 | -51 | -21 | 54 |
| L postcentral gyrus | BA 43 | cluster 1 | 7.87 | <.001 | -60 | -15 | 27 |
| L middle temporal gyrus | BA 21 | cluster 1 | 11.28 | <.001 | -54 | -45 | 30 |
| L middle temporal gyrus | BA 21 | cluster 1 | 6.88 | <.001 | -45 | -48 | 9 |
| R middle temporal gyrus | BA 20 | cluster 5 | 6.03 | <.005 | 69 | -42 | -6 |
| R middle temporal gyrus | BA 22 | cluster 5 | 4.86 | <.005 | 69 | -45 | 9 |
| R middle temporal gyrus | BA 21 | cluster 5 | 4.81 | <.005 | 54 | -33 | -6 |
| R superior parietal lobule | BA 7 | cluster 2 | 5.97 | <.001 | 15 | -63 | 51 |
| L inferior parietal lobule | BA 2 | cluster 1 | 12.62 | <.001 | -45 | -39 | 54 |
| L inferior parietal lobule | BA 3 | cluster 1 | 7.39 | <.001 | -54 | -24 | 39 |

| | | | | | | | |
|---|---|---|---|---|---|---|---|
| L inferior parietal lobule | BA 40 | cluster 1 | 13.06 | <.001 | -42 | -42 | 48 |
| L inferior parietal lobule | BA 40 | cluster 1 | 13.13 | <.001 | -36 | -45 | 45 |
| L inferior parietal lobule | BA 40 | cluster 1 | 13.10 | <.001 | -33 | -48 | 39 |
| R inferior parietal lobule | BA 2 | cluster 2 | 8.83 | <.001 | 51 | -36 | 51 |
| R angular gyrus | BA 40 | cluster 2 | 12.72 | <.001 | 33 | -51 | 39 |
| R supramarginal gyrus | BA 40 | cluster 2 | 9.13 | <.001 | 48 | -36 | 45 |
| R supramarginal gyrus | BA 40 | cluster 2 | 8.66 | <.001 | 39 | -33 | 39 |
| L cuneus | BA 18 | cluster 1 | 6.67 | <.001 | -6 | -84 | 24 |
| R cuneus | BA 18 | cluster 1 | 7.29 | <.001 | 6 | -78 | 21 |
| R cuneus | BA 17 | cluster 1 | 6.35 | <.001 | 15 | -96 | 12 |
| L middle occipital gyrus | BA 19 | cluster 1 | 6.73 | <.001 | -27 | -66 | 30 |
| R lingual gyrus | BA 18 | cluster 1 | 6.49 | <.001 | 9 | -75 | -3 |
| R lingual gyrus | BA 17 | cluster 2 | 5.80 | <.001 | 12 | -66 | 4 |
| R pallidum | | cluster 1 | 6.70 | <.001 | 12 | 0 | -3 |
| R putamen | | cluster 3 | 3.89 | <.001 | 30 | -3 | 6 |
| R cerebellum, lobule VI | | cluster 1 | 8.21 | <.001 | 30 | -66 | -24 |
| R cerebellum, crus I | | cluster 1 | 9.43 | <.001 | 39 | -60 | -27 |

Cluster volumes: 1=8810 voxels, 2=1170 voxels, 3=1075 voxels, 4=358 voxels, 5=139 voxels

**Supplementary table 4**

**Stereotaxic locations of peak voxels during discrimination (same sequences): all conditions – baseline**

| Brain area | Brodmann area | Cluster* | voxel t | pFDR (cluster) | x | y | z |
|---|---|---|---|---|---|---|---|
| L inferior frontal gyrus, p. opercularis | BA 44 | cluster 1 | 23.97 | <.001 | -45 | 6 | 27 |
| R inferior frontal gyrus, p. opercularis | BA 44 | cluster 1 | 28.42 | <.001 | 45 | 12 | 24 |
| R inferior frontal gyrus, p. triangularis | BA 45 | cluster 1 | 18.96 | <.001 | 51 | 21 | 9 |
| L anterior insula | BA 47 | cluster 1 | 35.15 | <.001 | -30 | 24 | -3 |
| R anterior insula | BA 47 | cluster 1 | 37.07 | <.001 | 36 | 21 | 0 |
| R anterior insula | BA 47 | cluster 1 | 36.93 | <.001 | 33 | 24 | -3 |
| L supplementary motor area | BA 6 | cluster 1 | 37.97 | <.001 | -3 | 0 | 60 |
| R supplementary motor area | BA 6 | cluster 1 | 31.58 | <.001 | 6 | 9 | 54 |
| L premotor cortex | BA 6 | cluster 1 | 19.32 | <.001 | -30 | -6 | 48 |
| L premotor cortex | BA 6 | cluster 1 | 25.90 | <.001 | -48 | -3 | 42 |
| R premotor cortex | BA 6 | cluster 1 | 28.38 | <.001 | 51 | 21 | 24 |
| R premotor cortex | BA 6 | cluster 1 | 27.09 | <.001 | 51 | 0 | 45 |
| L ventral premotor cortex | BA 6 | cluster 1 | 24.35 | <.001 | -51 | 6 | 15 |
| L ventral premotor cortex | BA 6 | cluster 1 | 23.93 | <.001 | -57 | 3 | 18 |
| L postcentral gyrus | BA 3 | cluster 1 | 20.05 | <.001 | -36 | -21 | 51 |
| L inferior parietal lobule | BA 40 | cluster 1 | 19.86 | <.001 | -39 | -39 | 45 |
| L inferior parietal lobule | BA 40 | cluster 1 | 18.16 | <.001 | -30 | -51 | 42 |
| R inferior parietal lobule | BA 40 | cluster 1 | 19.30 | <.001 | 33 | -51 | 42 |
| L superior temporal gyrus | BA 22 | cluster 1 | 66.06 | <.001 | -54 | -15 | 6 |
| L superior temporal gyrus | BA 41 | cluster 1 | 46.88 | <.001 | -45 | -36 | 12 |
| L superior temporal gyrus | BA 41 | cluster 1 | 44.01 | <.001 | -48 | -39 | 21 |
| L superior temporal gyrus | BA 41 | cluster 1 | 43.54 | <.001 | -54 | -39 | 18 |
| L superior temporal gyrus | BA 22 | cluster 1 | 40.03 | <.001 | -54 | -6 | -6 |
| L superior temporal gyrus | BA 22 | cluster 1 | 52.21 | <.001 | -66 | -30 | 12 |
| R superior temporal gyrus | BA 22 | cluster 1 | 57.44 | <.001 | 63 | -18 | 3 |
| R superior temporal gyrus | BA 22 | cluster 1 | 51.86 | <.001 | 63 | -27 | 9 |
| R superior temporal gyrus | BA 22 | cluster 1 | 47.61 | <.001 | 51 | -27 | 0 |
| L calcarine sulcus | BA 17 | cluster 1 | 39.96 | <.001 | -9 | -93 | 3 |
| R calcarine sulcus | BA 17 | cluster 1 | 38.94 | <.001 | 12 | -93 | 3 |
| R lingual gyrus | BA 18 | cluster 1 | 17.61 | <.001 | 21 | -72 | -6 |
| R fusiform | BA 19 | cluster 1 | 16.80 | <.001 | 30 | -66 | -9 |
| L thalamus | | cluster 1 | 16.76 | <.001 | -9 | -18 | 3 |

Cluster volumes: 1=22137 voxels

**Supplementary table 5**

**Stereotaxic locations of peak voxels during discrimination (same sequences): verbal condition – rhythm condition**

| Brain area | Brodmann area | Cluster* | voxel t | pFDR (cluster) | x | y | z |
|---|---|---|---|---|---|---|---|
| L inferior frontal gyrus, p. opercularis | BA 44 | cluster 3 | 3.91 | <.05 | -54 | 18 | 21 |
| L inferior frontal gyrus, p. triangularis | BA 44 | cluster 3 | 4.21 | <.05 | -42 | 15 | 27 |
| L superior temporal gyrus | BA 21 | cluster 1 | 5.98 | <.001 | -66 | -27 | 0 |
| L superior temporal gyrus | BA 22 | cluster 1 | 6.13 | <.001 | -60 | -15 | 6 |
| R superior temporal gyrus | BA 22 | cluster 2 | 5.25 | <.001 | 60 | -9 | -3 |
| L middle temporal gyrus | BA 22 | cluster 1 | 5.57 | <.001 | -63 | -36 | 3 |
| L middle temporal gyrus | BA 22 | cluster 1 | 4.82 | <.001 | -57 | -9 | -6 |
| R middle temporal gyrus | BA 21 | cluster 2 | 5.54 | <.001 | 66 | -21 | -3 |

Cluster volumes: 1=406 voxels, 2=259 voxels, 3=79 voxels

**Supplementary table 6**

**Stereotaxic locations of peak voxels during discrimination (different sequences): all conditions – baseline**

| Brain area | Brodmann area | Cluster* | voxel t | pFDR (cluster) | x | y | z |
|---|---|---|---|---|---|---|---|
| R middle frontal gyrus | BA 45 | cluster 1 | 18.23 | <.001 | 42 | 33 | 21 |
| L inferior frontal gyrus, p. triangularis | BA 44 | cluster 1 | 19.70 | <.001 | -54 | 21 | 12 |
| L inferior frontal gyrus, p. opercularis | BA 44 | cluster 1 | 24.90 | <.001 | -51 | 9 | 9 |
| R inferior frontal gyrus, p. opercularis | | cluster 1 | 35.88 | <.001 | 45 | 18 | 27 |
| L anterior insula | BA 47 | cluster 1 | 26.02 | <.001 | -30 | 24 | 0 |
| R anterior insula | BA 47 | cluster 1 | 30.83 | <.001 | 33 | 24 | 0 |
| L supplementary motor area | BA 6 | cluster 1 | 34.90 | <.001 | -6 | 6 | 54 |
| L supplementary motor area | BA 6 | cluster 1 | 37.45 | <.001 | -3 | 0 | 63 |
| R supplementary motor area | BA 6 | cluster 1 | 33.63 | <.001 | 6 | 12 | 51 |
| L premotor cortex | BA 6 | cluster 1 | 27.59 | <.001 | -54 | 0 | 45 |
| L premotor cortex | BA 6 | cluster 1 | 19.42 | <.001 | -33 | -15 | 60 |
| L premotor cortex | BA 6 | cluster 1 | 23.08 | <.001 | -45 | -3 | 54 |
| R premotor cortex | BA 6 | cluster 1 | 29.49 | <.001 | 51 | 0 | 45 |
| L ventral premotor cortex | BA 6 | cluster 1 | 21.42 | <.001 | -45 | 3 | 27 |
| L ventral premotor cortex | BA 6 | cluster 1 | 22.47 | <.001 | -57 | 6 | 18 |
| L superior temporal gyrus | BA 41 | cluster 1 | 39.69 | <.001 | -42 | -39 | 18 |
| L superior temporal gyrus | BA 22 | cluster 1 | 59.82 | <.001 | -63 | -24 | 6 |
| L superior temporal gyrus | BA 22 | cluster 1 | 58.40 | <.001 | -54 | -24 | 6 |
| L superior temporal gyrus | BA 22 | cluster 1 | 53.91 | <.001 | -60 | -15 | 3 |
| R superior temporal gyrus | BA 22 | cluster 1 | 61.46 | <.001 | 60 | -27 | 6 |
| R superior temporal gyrus | BA 22 | cluster 1 | 51.26 | <.001 | 57 | -36 | 9 |
| R superior temporal gyrus | BA 22 | cluster 1 | 45.89 | <.001 | 51 | -15 | -3 |
| L postcentral gyrus | BA 4 | cluster 1 | 24.52 | <.001 | -36 | -27 | 51 |
| R inferior parietal lobule | BA 2 | cluster 1 | 24.10 | <.001 | 48 | -36 | 48 |
| R supramarginal gyrus | BA 40 | cluster 1 | 21.57 | <.001 | 45 | -39 | 45 |
| L inferior parietal lobule | BA 40 | cluster 1 | 19.33 | <.001 | -39 | -42 | 45 |
| R inferior parietal lobule | BA 40 | cluster 1 | 20.46 | <.001 | 36 | -51 | 45 |
| R inferior parietal lobule | BA 2 | cluster 1 | 19.63 | <.001 | 51 | -36 | 57 |
| R cuneus | BA 17 | cluster 1 | 26.90 | <.001 | 15 | -99 | 15 |
| L calcarine sulcus | BA 17 | cluster 1 | 43.53 | <.001 | -9 | -96 | 3 |
| R calcarine sulcus | BA 17 | cluster 1 | 33.17 | <.001 | 6 | -90 | 0 |
| L inferior occipital gyrus | BA 19 | cluster 1 | 18.74 | <.001 | -27 | -81 | -9 |

Cluster volumes: 1=21626 voxels

**Supplementary table 7**

**Stereotaxic locations of peak voxels during discrimination (different sequences): verbal condition – rhythm condition**

| Brain area | Brodmann area | Cluster* | voxel t | pFDR (cluster) | x | y | z |
|---|---|---|---|---|---|---|---|
| L superior frontal gyrus | BA 9 | cluster 4 | 4.69 | <.001 | -18 | 45 | 39 |
| R superior frontal gyrus | BA 9 | cluster 3 | 5.67 | <.001 | 15 | 48 | 39 |
| R superior frontal gyrus | BA 9 | cluster 3 | 4.90 | <.001 | 18 | 39 | 45 |
| L medial superior frontal gyrus | BA 9 | cluster 4 | 4.73 | <.001 | -9 | 54 | 33 |
| Medial superior frontal gyrus | BA 10 | cluster 5 | 3.60 | <.005 | 0 | 54 | 9 |
| L middle frontal gyrus | BA 8 | cluster 4 | 5.54 | <.001 | -30 | 18 | 51 |
| L middle frontal gyrus | BA 8 | cluster 4 | 5.39 | <.001 | -24 | 24 | 54 |
| L middle frontal gyrus | BA 9 | cluster 4 | 5.01 | <.001 | -27 | 30 | 48 |
| R middle frontal gyrus | BA 8 | cluster 3 | 6.68 | <.001 | 27 | 27 | 48 |
| L inferior frontal gyrus, p. opercularis | BA 44 | cluster 4 | 4.16 | <.001 | -51 | 18 | 36 |
| R medial orbitofrontal cortex | BA 11 | cluster 5 | 4.82 | <.005 | 6 | 45 | -12 |
| L superior parietal lobule | BA 7 | cluster 2 | 4.40 | <.001 | -24 | -54 | 42 |
| L inferior parietal lobule | BA 40 | cluster 2 | 4.92 | <.001 | -54 | -48 | 39 |
| R supramarginal gyrus | BA 40 | cluster 1 | 6.38 | <.001 | 57 | -48 | 36 |
| L angular gyrus | BA 39 | cluster 2 | 5.68 | <.001 | -45 | -51 | 33 |
| R angular gyrus | BA 39 | cluster 1 | 6.75 | <.001 | 51 | -63 | 33 |
| R angular gyrus | BA 39 | cluster 1 | 6.59 | <.001 | 54 | -60 | 24 |
| R angular gyrus | BA 7 | cluster 1 | 5.80 | <.001 | 39 | -69 | 42 |
| L middle cingulate gyrus | BA 23 | cluster 2 | 6.55 | <.001 | -6 | -45 | 36 |
| L middle cingulate gyrus | BA 23 | cluster 2 | 6.29 | <.001 | -3 | -36 | 39 |
| L middle cingulate gyrus | BA 23 | cluster 2 | 5.83 | <.001 | -3 | -69 | 33 |
| R middle cingulate gyrus | BA 23 | cluster 2 | 7.64 | <.001 | 12 | -42 | 36 |
| R middle cingulate gyrus | BA 23 | cluster 2 | 7.06 | <.001 | 6 | -33 | 33 |
| R posterior cingulate gyrus | BA 26 | cluster 2 | 6.18 | <.001 | 3 | -45 | 24 |
| R precuneus | BA 7 | cluster 2 | 6.02 | <.001 | 6 | -69 | 39 |
| R precuneus | | cluster 2 | 7.39 | <.001 | 9 | -51 | 45 |
| L superior temporal gyrus | BA 22 | cluster 2 | 7.21 | <.001 | -63 | -21 | 3 |
| L middle temporal gyrus | BA 21 | cluster 2 | 7.01 | <.001 | -66 | -39 | 0 |
| L middle temporal gyrus | BA 21 | cluster 2 | 6.10 | <.001 | -57 | -24 | -6 |
| L middle temporal gyrus | BA 20 | cluster 2 | 6.03 | <.001 | -54 | -27 | -9 |
| L middle temporal gyrus | BA 22 | cluster 2 | 6.33 | <.001 | -60 | -12 | -3 |
| L middle temporal gyrus | BA 39 | cluster 2 | 6.82 | <.001 | -48 | -63 | 18 |
| L middle temporal gyrus | BA 20 | cluster 2 | 4.27 | <.001 | -57 | -42 | -15 |
| L middle temporal gyrus | BA 37 | cluster 2 | 5.51 | <.001 | -63 | -51 | 24 |
| L middle temporal gyrus | BA 37 | cluster 2 | 4.66 | <.001 | -48 | -57 | -12 |
| R middle temporal gyrus | BA 31 | cluster 2 | 6.31 | <.001 | -63 | -48 | 3 |
| R middle temporal gyrus | BA 22 | cluster 1 | 8.50 | <.001 | 60 | -15 | -9 |
| R middle temporal gyrus | BA 21 | cluster 1 | 7.53 | <.001 | 63 | -24 | -3 |
| R middle temporal gyrus | BA 37 | cluster 1 | 6.69 | <.001 | 45 | -66 | 27 |
| R middle temporal gyrus | BA 31 | cluster 1 | 5.07 | <.001 | 63 | -42 | 0 |
| L inferior temporal gyrus | BA 20 | cluster 2 | 4.33 | <.001 | -57 | -48 | -15 |
| L inferior temporal gyrus | BA 20 | cluster 2 | 5.55 | <.001 | -63 | -21 | -21 |
| L inferior temporal gyrus | BA 37 | cluster 2 | 5.40 | <.001 | -51 | -51 | -6 |
| L inferior temporal gyrus | BA 37 | cluster 2 | 4.41 | <.001 | -57 | -63 | -6 |

| | | | | | | | |
|---|---|---|---|---|---|---|---|
| R inferior temporal gyrus | BA 20 | cluster 1 | 5.19 | <.001 | 63 | -15 | -24 |
| R middle occipital gyrus | BA 19 | cluster 1 | 6.25 | <.001 | 36 | -72 | 33 |
| L middle occipital gyrus | BA 19 | cluster 2 | 7.12 | <.001 | -39 | -72 | 33 |
| L cuneus | | cluster 2 | 4.74 | <.001 | -15 | -69 | 27 |
| L superior occipital gyrus | BA 19 | cluster 2 | 4.34 | <.001 | -21 | -63 | 30 |
| L fusiform gyrus | BA 37 | cluster 2 | 4.22 | <.001 | -45 | -57 | -21 |
| L cerebellum lobule VI | | cluster 2 | 4.24 | <.001 | -42 | -54 | -24 |

Cluster volumes: 1=1982 voxels, 2=4428 voxels, 3=513 voxels, 4=510 voxels, 5=198 voxels

### Supplementary table 8

**Stereotaxic coordinates for region-of-interest (ROI) analyses**

| **Encoding** | | | | **Maintenance** | | | | **Discrimination** | **(same)** | | | **(different)** | | |
|---|---|---|---|---|---|---|---|---|---|---|---|---|---|---|
| **Region** | **x** | **y** | **z** | **Region** | **x** | **y** | **z** | **Region** | **x** | **y** | **z** | **x** | **y** | **z** |
| SMA | -6 | 3 | 60 | L insula | -42 | 15 | 6 | L PMC | -48 | -3 | 42 | -54 | 0 | 45 |
| L dPMC | -51 | 0 | 45 | L prefrontal cortex | -42 | 30 | 30 | L parietal cortex | -39 | -39 | 45 | -39 | -42 | 45 |
| R dPMC | 51 | 3 | 45 | L vPMC | -51 | 3 | 42 | L STG | -54 | -15 | 6 | -63 | -24 | 61 |
| L putamen | -21 | 6 | 3 | L anterior insula | -30 | 24 | 0 | L anterior insula | -30 | 24 | 3 | -30 | 24 | 0 |
| R putamen | 21 | 6 | 0 | L dorsomedial PMC | -33 | -66 | 3 | L occipital cortex | -9 | -93 | 3 | -9 | -93 | 6 |
| L STG 1 | -54 | -21 | 6 | L parietal cortex | -33 | -48 | 39 | L vPMC | -51 | 6 | 15 | -51 | 9 | 9 |
| L STG 2 | -66 | -21 | 6 | R prefrontal cortex | 36 | 39 | 24 | R PMC | 51 | 0 | 45 | 51 | 0 | 45 |
| R STG 1 | 57 | -18 | 0 | R anterior insula | 33 | 24 | -3 | R parietal cortex | 33 | -51 | 42 | 48 | -36 | 48 |
| R STG 2 | 63 | -3 | 0 | R cerebellum | 39 | -60 | -27 | R STG | 63 | -18 | 3 | 60 | -27 | 6 |
| L anterior insula | -36 | 21 | 3 | R parietal cortex | 33 | -51 | 39 | R anterior insula | 36 | 21 | 0 | 33 | 24 | 0 |
| R anterior insula | 33 | 21 | 3 | SMA | -3 | 6 | 54 | R occipital cortex | 12 | -93 | 3 | 6 | -90 | 0 |
| L occipital | -9 | -93 | -3 | | | | | R vPMC | 45 | 12 | 24 | 45 | 18 | 27 |
| R occipital | 12 | -96 | 3 | | | | | SMA | -3 | 0 | 60 | -3 | 0 | 63 |
| L vPMC | -51 | 9 | 6 | | | | | | | | | | | |
| R vPMC | 45 | 6 | 27 | | | | | | | | | | | |
| R parietal | 33 | -51 | 45 | | | | | | | | | | | |
| L cerebellum | -27 | -63 | -21 | | | | | | | | | | | |
| R cerebellum | 30 | -63 | -21 | | | | | | | | | | | |
